# Locus coeruleus norepinephrine selectively controls visual attention

**DOI:** 10.1101/2022.09.30.510394

**Authors:** Supriya Ghosh, John H. R. Maunsell

## Abstract

Norepinephrine (NE) neuromodulation plays a role in diverse non-specific physiological and cognitive functions including wakefulness, arousal, and cognitive performance. NE modulation of neuronal responses in the cerebral cortex has been proposed to mediate improved task-specific behavior by enhancing sensory processing. However, the sensory-specific NE contribution on performance remains unknown. We directly tested the role of NE-mediated neuromodulation of sensory signals on perceptual performance in non-human primates doing visual spatial attention tasks. We found that NE neurons in the locus coeruleus (LC) respond selectively to an attended stimulus. Optogenetically enhancing the sensory-specific responses of LC-NE neurons improved the monkeys’ sensory discrimination in a spatially selectively way, without affecting motor processing. These findings identify a specific contribution of NE neuromodulation of sensory representations to selective attention and performance.

**One-Sentence Summary:** Optogenetic activation of monkey locus coeruleus causes a strong and spatially selective improvement in visual sensitivity.

**Graphical abstract:** Elevation of phasic norepinephrine signal by optogenetic stimulation of locus coeruleus in non-human primates selectively improves attentional performance that attributes to enhanced sensory sensitivity.

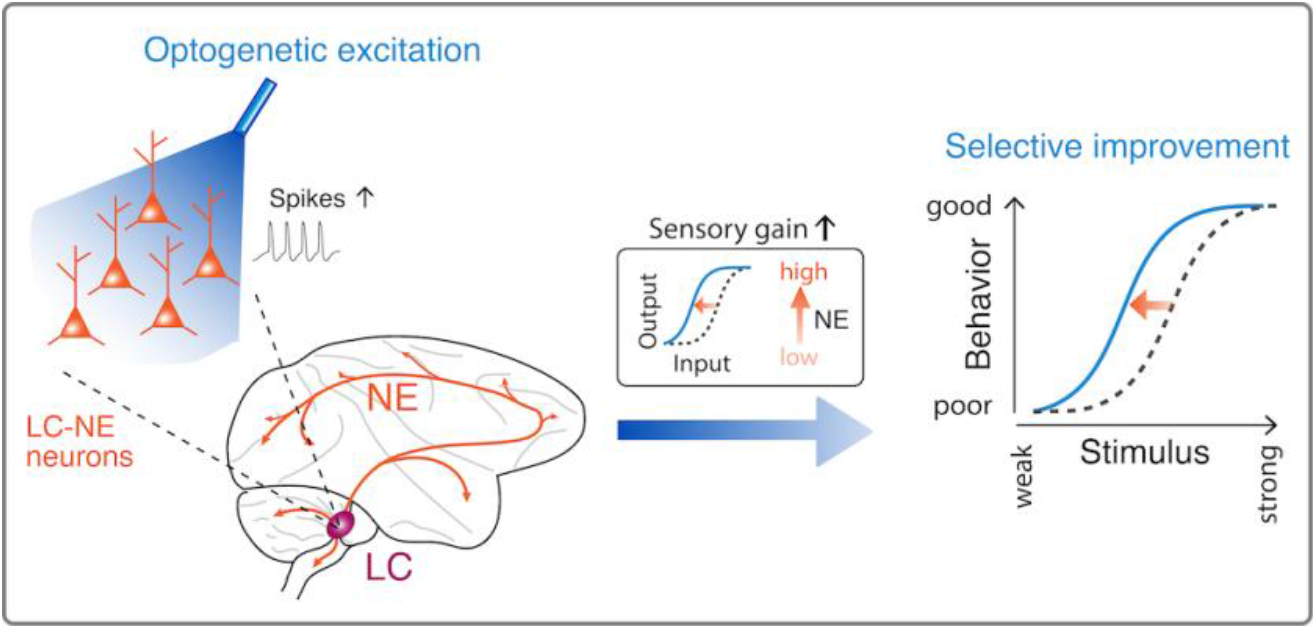

## Main Text

Neuromodulators have long been thought to play crucial roles in task specific cognitive functions, including selective attention (*1*). Neuromodulation by norepinephrine (NE) has a powerful effect on general arousal and wakefulness (*2*). The locus coeruleus (LC) is a primary source of NE in the brain and is known to contribute to diffuse, non-specific arousal (*3–5*). However, a correlation between LC activation and behavioral accuracy in non-human primates (*6*) coupled with NE-induced enhancement of signal-to-noise in sensory representations in the cerebral cortex (*3, 7, 8*) suggest that LC-NE might improve perceptual performance by selectively facilitating relevant sensory inputs, thereby contributing to selective attention. It remains unknown whether NE-mediated effects on sensory processing have a causal role on attentional performance in primates, owing to previous limitations on temporally-precise control of LC-NE activity in non-human primates doing tasks that allow control of attention and measurement of sensory performance free of confounds from motor actions.

We therefore designed a visual task to control monkeys’ behavioral performance that selectively relied on sensory sensitivity independent of decision and motor selection. We measured neuronal responses from LC-NE neurons and optogenetically enhanced their responses to task-relevant visual stimuli to measure their contributions to spatially selective attentional performance. We found that LC neurons convey spatially selective signals related to visual attention. By optogenetically activating these LC neurons, we identified specific causal roles of stimulus-locked neuronal activity of LC-NE neurons on improving behavioral performance to attended visual stimuli.

We trained two rhesus monkeys to selectively attend one of two visual stimulus locations while doing a visual orientation change detection task (**Fig. 1A**) (*9*). Once the animal fixated, two Gabor sample stimuli appeared for 200 ms, one in each hemifield. After a brief delay (200-300 ms), a Gabor test stimulus appeared in one of the locations. The monkey had to report whether the test stimulus had a different orientation than the sample that had appeared in that location by making a saccade to one of two colored saccade targets after the fixation spot went off. The locations of the saccade targets were interchanged randomly across trials. In this way, signals related to motor planning (*10*) were dissociated from the attention-related perceptual sensitivity we wanted to examine.

**Figure 1.**
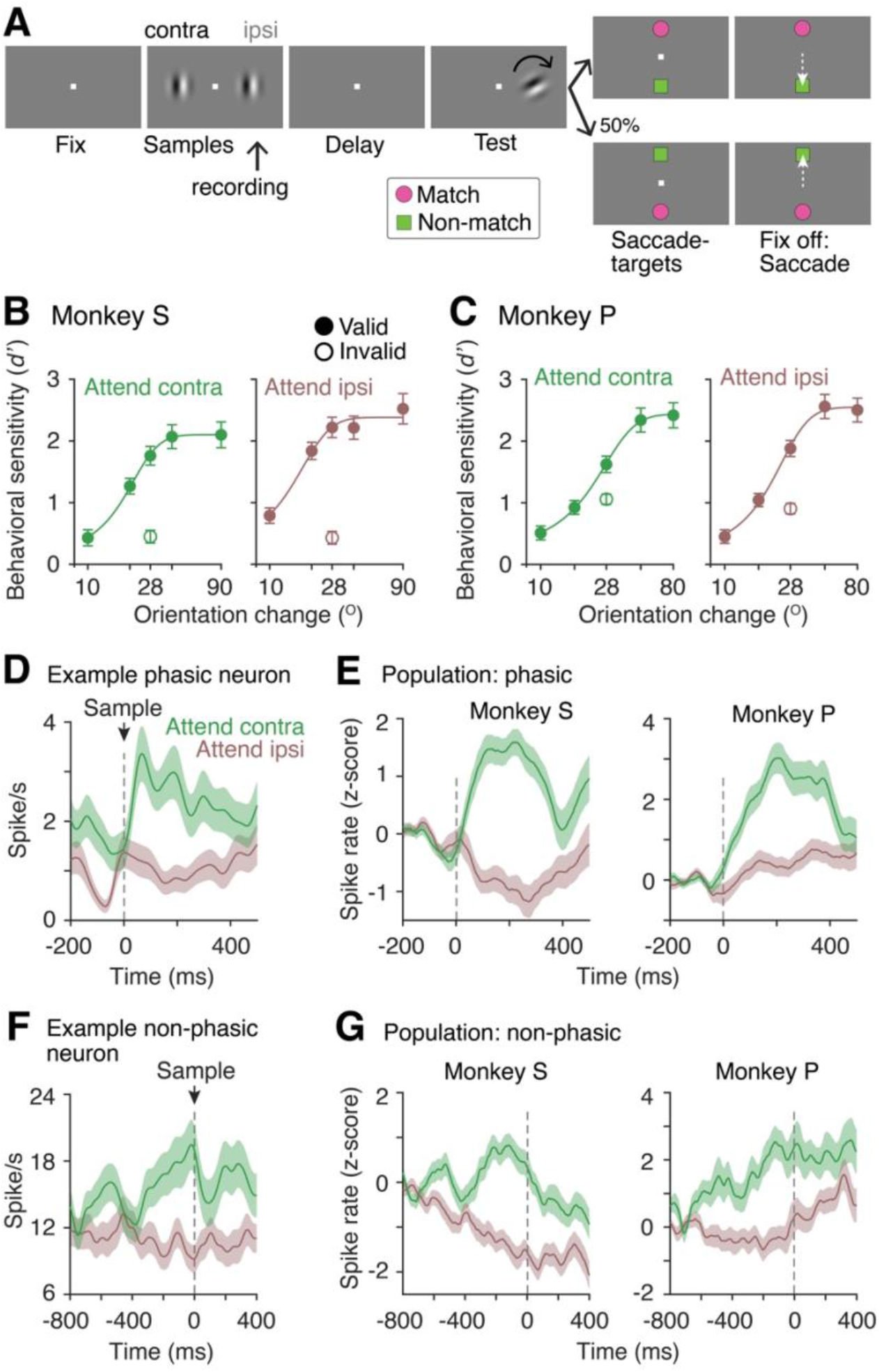
Modulation of LC neurons to spatially selective visual attention. (**A**) Visual orientation change detection task. Monkeys maintained fixation (400–800 ms) while attending to one of the two sample Gabor presented (200 ms) in opposite hemifields. After a short delay (200–300 ms), a test stimulus (200 ms) appeared at one of the stimuli locations. Monkeys reported an orientation change between sample and test by making a saccade to the appropriate saccade-target (different colors and shapes). Positions of the saccade targets were randomized on each trial. Monkeys’ spatial attention was controlled between the stimuli locations in alternate blocks of trials by varying the reward size for correct detections. (**B-C**) Monkeys’ performance averaged across sessions as a function of orientation changes (#sessions, monkey S, 29; monkey P, 22). Behavioral sensitivity (*d’*) is much better on trials when the test stimulus appeared at the attended location (valid trials; filled circles) compared to the unattended location (invalid trials; open circles). Error bars, 95% confidence intervals. (**D-E**) Example and population PSTHs of spike rates of phasic LC neurons for all correct trials when monkeys attended either contralateral or ipsilateral to recorded LC (phasic, monkey S, n = 89, monkey P, n = 74; non-phasic, monkey S, n = 88, monkey P, n = 53). Error bars, ± 1 SEM. (**F-G**) Same as in (D-E), but for non-phasic LC neurons.

Spatially selective attention substantially improved monkeys’ detection performance at the attended location relative to the unattended location, measured by behavioral sensitivity (*d*’) (**Fig. 1B-C**). While the monkeys did the attention task, we recorded simultaneously from populations of LC neurons (monkey S, n = 229; monkey P, n = 185). According to the responses to task events, recorded neurons were classified as phasic responsive to visual stimulus (phasic; monkey S, n =), pre-stimulus fixation responsive (non-phasic) and saccade responsive (saccade) (**Supplementary fig. 3**) Responses of phasic LC neurons to the sample stimuli were stronger when attention was directed to the stimulus contralateral to the recorded LC (**Fig. 1D-E**; Monkey S, neuronal modulation (*d*’_neuron_) over sample period = 0.84 ± 0.09, p < 10^-15^; Monkey P, *d’*_neuron_ = 0.65 ± 0.07, p < 10^-9^; signed-rank test). Selective attention also increased the spiking of non-phasic LC neurons (**Fig. 1F-G**; Monkey S, *d*’_neuron_ over the fixation period = 0.78 ± 0.05, p < 10^-15^; Monkey P, *d*’_neuron_ = 0.67 ± 0.06, p < 10^-13^; signed-rank test). Responses of saccade neurons were unchanged throughout the trial (**Supplementary fig. 4**). This modulation of LC activity by attentional shifts was associated with an increased behavioral *d*’, and was independent of perceptual decision criterion and motor criterion (**Supplementary fig. S1**).

Previous studies suggest that the spiking of LC neurons closely varies with behavioral performance (*6*). Thus, we decomposed population firing rates into demixed principal components (dPC) (*11*) to quantify how activity of LC neurons relates to perceptual detection (correct versus error) and the focus of selective attention (contra versus ipsi hemifields relative to the recorded LC) in single trials (**Materials and Methods; Fig. 2A, 2C; Supplementary figs. S5, S6**). Spike trains of LC neurons carried significant information about selective attention and detection of stimulus orientation change as measured by decoding accuracy in cross-validated (leave-one-out) single trials (**Fig. 2B, 2D**).

**Figure 2.**
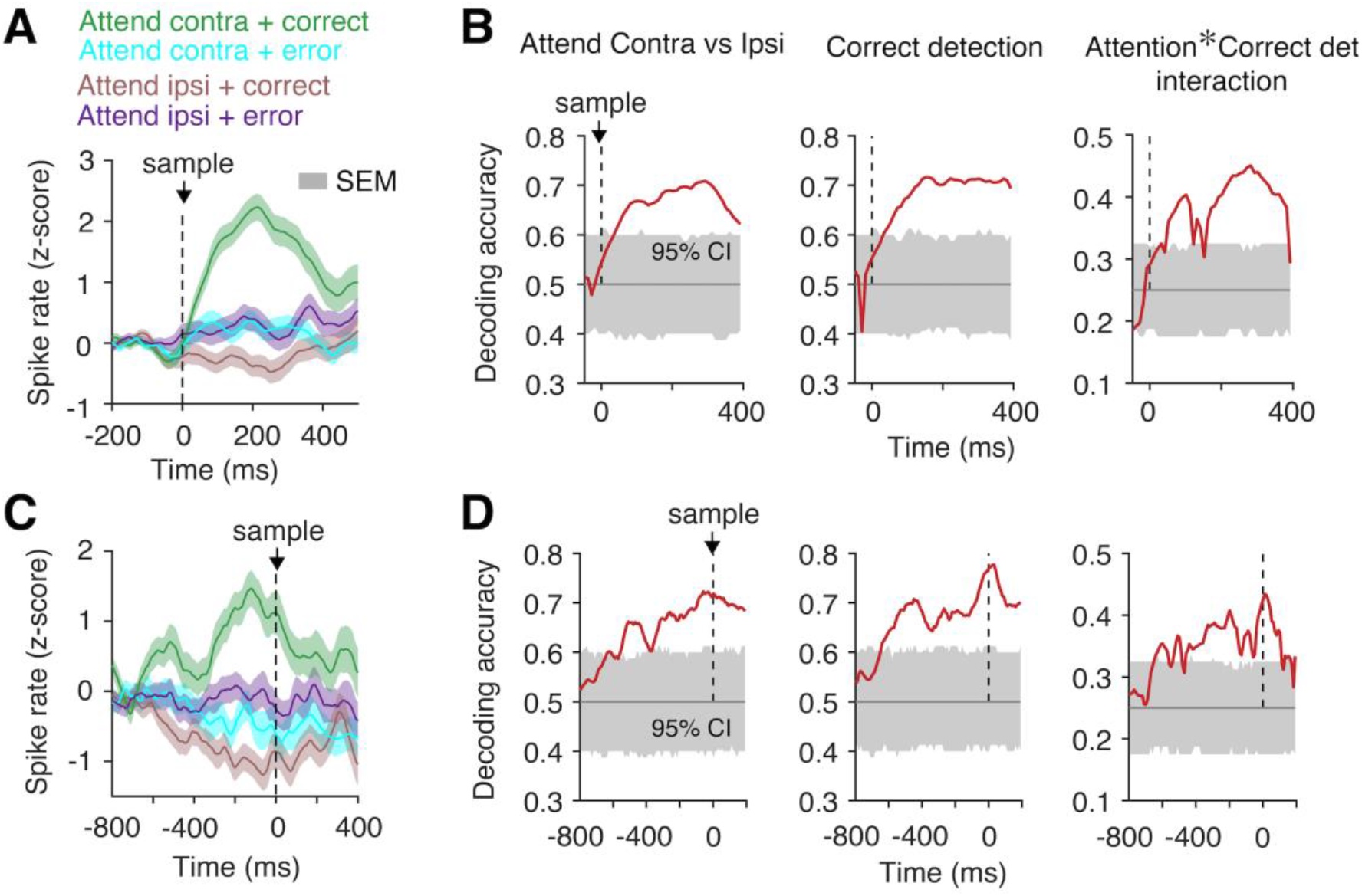
Single trial prediction of selective attention and sensory detection by LC population code. (**A**) Population PSTHs spike rates of phasic LC neurons for correct and error trials while attention was directed either contra- or ipsi-lateral to the recorded LC (n = 163, two monkeys). Error bars, ± 1 SEM. (**B**) Time course of single-trial decoding accuracy (cross-validation leave-one out trials) for different cognitive variables: selective attention, correct detection and their interaction based on demixed principal components of population PSTHs of phasic LC neurons as in (A) (**Material and Methods**). Error bars, 95% confidence intervals from shuffled trials (**C-D**) Same as in (A-B), but for non-phasic LC neurons (n = 141, two monkeys).

Further, some dPCs represented significant information about the attention-by-detection interaction (**Supplementary figs. S5, S6; Fig. 2B, 2D**). These components isolated the focus of trial-by-trial enhanced sensory sensitivity-attentional engagement that directly relate to correctly detecting a stimulus change. Similar to the LC, neurons in other neuromodulatory systems including dopamine and acetylcholine are also known to exhibit mixed selectivity to multiple task relevant factors (*1*).

To directly examine causal roles of stimulus-locked attentional modulation of LC-NE neurons, we selectively expressed excitatory channelrhodopsin (ChR2) in NE neurons in the LC. The same monkeys were unilaterally injected with two adeno-associated viruses (**Materials and Methods**) in an approach that has been found to give robust cell-type specific opsin expression in monkeys (*12*) (**Supplementary fig. S7**). Subsequently, NE neurons in the LC were identified by characteristic neurophysiological responses to salient sound stimuli (*13, 14*) and excitation by optical stimulation using brief blue light (**Fig. 3A, 3B**).

**Figure 3.**
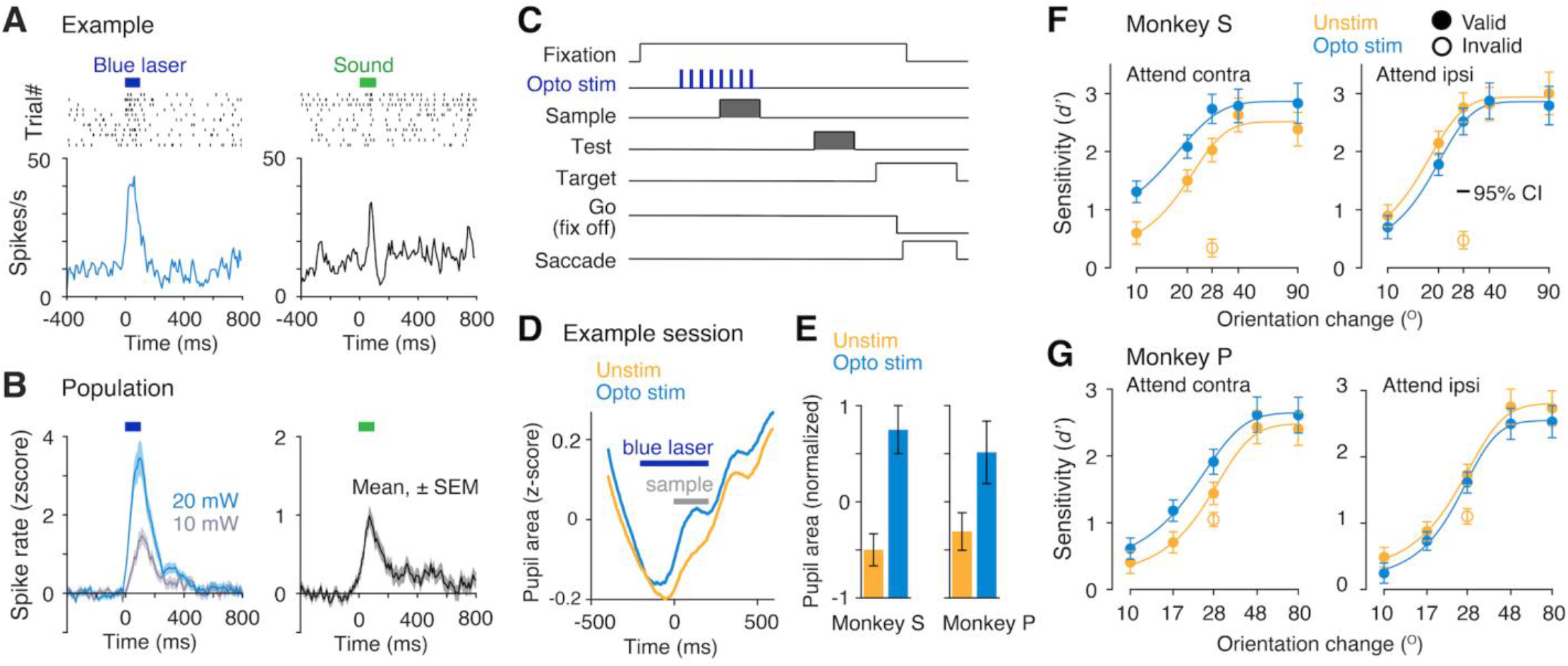
Optogenetic excitation of LC-NE neurons improves attentional performance contralateral to the stimulated LC. (**A**) Example LC-NE neuron. *Left*, Optogenetic excitation by brief (100 ms) blue light laser pulses (*top*, single trial spike trains; *bottom*, spike rate PSTH). *Right*, response to brief white-noise auditory sound. (**B**) Same as in (A), for the population of same LC-NE neurons in Fig. 1. Optogenetic stimulation was tested for two laser intensities, 10 and 20 mW. (**C**) Optogenetic stimulation during the sample stimulus in spatially selective visual attention task (Fig. 1A). LC was stimulated unilaterally on a randomly interleaved 50% of valid trials by blue laser pulses over 400 ms, at 20 Hz, 10 ms pulse width, 40 mW power, starting 200 ms before sample stimulus onset. (**D**) Pupil area in an example session aligned with respect to sample onset. Optogenetic stimulation enhanced pupil area compared to unstimulated trials. (**E**) Session averaged mean pupil area for unstimulated and LC stimulated trials for monkeys S and P. Error bars, ± 1 SEM. (**F-G**) Session averaged behavioral sensitivities (*d’*) of monkey S (F; n = 11) and monkey P (G; n = 11) as a function of orientation changes. LC activation improved *d’s* compared to unstimulated trials when animal’s attention was directed to the stimulus contralateral to the stimulated LC. Error bars, 95% confidence intervals.

On a random half of the trials, we optogenetically activated LC-NE neurons while the monkeys did the attention task (**Fig. 3C**). These stimulation sessions were a subset of the total experimental sessions used for the neurophysiological recordings in **Figs. 1** and **2**. A 400 ms train of eight 10 ms optical pulses (20 Hz) was delivered to the LC starting 200 ms before the sample stimuli. Previous studies suggest that the neuronal activity of the LC-NE system is closely linked with the pupillary area (*15*), which provides an index of arousal and cognitive engagement. Unilateral optogenetic activation of LC-NE neurons increased pupil area relative to the unstimulated trials, documenting the efficacy of the LC-NE stimulation to alter physiological state (**Fig. 3D, 3E**, monkey S, p < 0.01; monkey P, p < 0.01; paired *t*-test). When the animal’s attention was directed to the stimulus in the visual hemifield contralateral to the stimulated LC, detection of orientation changes in that hemifield was greatly enhanced (blue curve, **Fig. 3F, 3G**), with no significant increase in decision criterion (**Supplementary fig. S2**). In contrast, LC activation caused a weak impairment of the animal’s discrimination of ipsilateral orientation changes. Thus, LC activity can direct spatially-targeted visual attention. Further, a uniform enhancing effect of LC stimulation on pupil area irrespective of the spatial focus of monkeys’ attention (monkey S, p = 0.94; monkey P, p = 0.86; paired *t*-test) suggest that LC-NE mediated selective effect on performance was not driven by enhanced effective retinal illumination due to larger pupil area.

This spatially selective effect of LC stimulation was not driven by enhanced effective retinal illumination due to larger pupil area (**Fig. 3D, 3E**). LC-stimulation uniformly affected pupil area irrespective of the location of monkeys’ spatial attention.

Previous studies have reported bilateral or non-specific effects of optogenetic unilateral LC activation or inhibition on behavior in rodents (*5, 16*). Thus, we tested whether this specific effect of LC activation on performance in our task depended on the selectivity of spatial attention. In separate experimental sessions, we varied the animals’ spatially non-selective attention (attentional effort) between low and high values with attention directed equally to both hemifields (**Fig. 4**). We optogenetically stimulated LC-NE neurons unilaterally on random half of the trials. The monkeys’ detection performance improved selectively in the contralateral location without any effects on the ipsilateral detection. Together, these results on LC stimulation during selective attention and non-selective attentional effort confirm that LC-NE activation can drive a spatially selective enhancing effect on perceptual performance to the behaviorally relevant stimulus.

**Figure 4.**
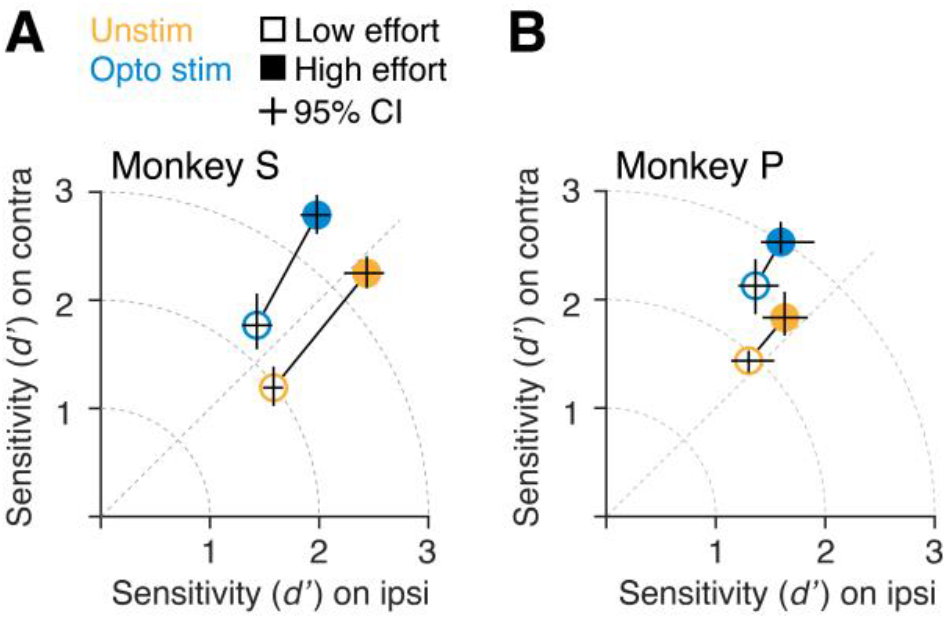
Effects of LC-NE activation on behavioral performance is independent of attentional selectivity. (**A-B**) Session averaged behavioral *d’*s for contra- and ipsi-lateral stimuli for monkey S (A; n = 9) and monkey P (B; n = 9) when animal’s attention was equally distributed between the two hemifields (non-selective attentional effort; yellow circles) using the visual task described in Figure 1A. Attentional effort was controlled between low and high values between alternate blocks of trials by varying reward size for correct detections. Optogenetic stimulation (randomly interleaved 50% of trials; blue circles) during the sample stimulus selectively improved behavioral *d’* to the contralateral stimulus. Error bars, 95% confidence intervals.

In agreement with other studies (*10, 13, 14*), our results suggest that LC neurons are responsive to different task relevant stimuli and behavioral responses. Increased phasic LC spiking associated with rewarded response target (*13*) or effortful motor action (*10*) has been shown to be associated with attentional orientation toward a relevant response. Here, we additionally isolated a distinct spike rate modulation in the LC that is associated with increased visual spatial attention to a contralateral location. Improved behavioral performance in our task at the attended location was associated with increased sensory sensitivity, and was independent of perceptual decision and motor response. This observed LC spike rate modulation closely relates to increased sensory sensitivity or sensory gain seen in many cortical and subcortical visual areas associated with increased behavioral sensitivity in the neuron’s response field (*9, 17, 18*). Notably, attention-related modulations in LC approach an all-or-none effect depending on behavioral relevance of the visual stimulus, and are far greater than typical attentional modulations seen in visual cerebral cortex (*9, 17*). However, a direct estimate of the amount of NE release in the primate visual cortex associated to an attended sensory stimulus would be valuable to better understand the complexities of local computations (*19, 20*). Although, synchronous activation of population of LC-NE neurons in attention demanding tasks might elicit large number of spikes collectively (*6*), the precise levels of NE changes in distinct cortical areas owing to changes in few spikes per LC neuron (8.1 ± 0.4 spike/s, mean and ± 1 SEM) in perceptual tasks like ours need to be investigated.

To the best of our knowledge, the consequences of sensory-evoked phasic NE modulation on task-specific sensory signals and behavior have not been investigated previously. The strongly coupled activity of phasic and tonic spiking of LC neurons (*6*) has limited possibilities for establishing the roles of LC activation on task-specific sensory processing. Previous studies showed that pharmacological enhancement or blocking of central NE release improves or impairs target-specific responsivity, respectively, affecting visual attention performance in monkeys (*21*) and humans (*22*). Electrical stimulation of LC was found to improve memory retention in rats (*23*). Recent cell-type specific optogenetic activation of LC-NE neurons in rodents produced more general arousal (sleep to-awake transition) (*5*) or motor response-specific effects (*24*). These earlier studies targeting NE system using pharmacology, electrical stimulation or optogenetics lacked precise control on temporal specificity, cell-type specificity and task-specific cognitive function e.g., sensory-versus-response selectivity during attention tasks. Our results show that temporally restricted phasic activation of specifically NE neurons during the sensory selection is sufficient to improve performance to contralateral visual stimulus in spatial attention task. This identifies a distinct contribution of NE activity in mediating task-relevant selective sensory processing and associated behavior. Combining activation of LC-NE neurons with recordings from visual structures would help to further illuminate the role of LC-NE in attentional control.

One important question is the mechanisms that support NE-mediated spatially selective sensory selection in visual areas targeted by the LC. Our result on unilateral attentional effects of LC optogenetic stimulation in monkeys is surprising, given the widespread LC projections throughout the cerebral cortex, and moderate but non-specific (bilateral) effects of unilateral LC stimulation on the contralateral LC and prefrontal cortex (PFC) in rodents (*25*). However, LC-NE neuronal spike modulation with selective attention and unilateral effect of optogenetic stimulation are consistent with the LC’s dominant ipsilateral connectivity with the forebrain and visual cortex (*26*), and neurophysiological evidence that phasic LC discharge rapidly increases the signal-to-noise of sensory representations in the cerebral cortex (*3, 8*). Further, activation of this phasic LC response can mimic activity in the sensory cortex and perception of a salient stimulus (*27*), given the strong influence of neuromodulators in early visual processing (*28*).

Spatially selective attention is known to strongly modulate neuronal responses in many brain areas in the visual system (*29*) that receive dense LC-NE projections, including dorsolateral PFC, frontal eye field, area V4, and superior colliculus. Our results, combined with empirical evidence on stimulus-specific selective effects of localized NE in the visual cortex (*8*), support the previously proposed “glutamate amplifies noradrenergic effects” (GANE) model (*30*) of spatially selective attentional spike modulation in the visual system. In this model, high glutamate release in response to the attended stimulus creates a localized NE hotspot via positive feedback between the glutamate and NE release. Such increased NE concentration could amplify the salient stimulus evoked response further. In contrast, NE depletion would result in a suppressive effect for representations of unattended stimuli. Detailed sensory specific distribution of LC-NE projections in multi-sensory areas are poorly characterized. Thus, our findings in addition to anatomical mapping of sensory-specific selective LC activation would be crucial for better understanding of distinct neuromodulatory controls on perception and cognition.

Altogether, this study provides important experimental evidence for a distinct NE neuromodulatory role in primates that regulates sensory sensitivity to improve attention-demanding perceptual performance in a spatially selective way. The findings have potential clinical implications for designing targeted therapeutic interventions according to the specific deficits in sensory, decision or motor performance observed in disorders of attention, including attention-deficit/hyperactivity disorder.

## Acknowledgments

We thank Dr. Marlene R. Cohen, Dr. Anita A. Disney, Chery J. Cherian and Lai Wei for critical feedback on the manuscript; Dr. Jackson J. Cone and Dr. Mitchell F. Roitman for assistance with optrode fabrication; Morgan L. Bade, Rachel Parker and Autumn O. Mitchell for technical help with monkey procedures.

## Author contributions

S.G. and J.H.R.M. designed the experiments, performed the surgeries, and wrote the paper. S.G. performed the experiments and analyzed the data.

## Data and materials availability

All data are available in the main text or the supplementary materials (external Data S1-S2).

## Supplementary Materials

Materials and Methods

Supplementary Text

Figs. S1 to S7

References (*1–32*)

