## Supplementary Materials for "Locus coeruleus norepinephrine selectively controls visual attention"

**This PDF file includes:**

Materials and Methods
Supplementary Text
Figs. S1 to S7

### Materials and Methods

#### Animals and surgery

Two adult male rhesus monkeys (*Macaca mulatta*, 9 and 13 kg) were implanted with a titanium head post using aseptic surgical techniques before behavioral training began. After the completion of training (3 to 5 months), we surgically implanted a stainless-steel recording chamber targeting the LC on one side (monkey S, right; monkey P, left) to access the LC, guided by an MRI obtained before the initial surgery. The cylinders were centered on the skull at intra-aural zero coordinates 3.5 mm A, 10.0 mm L, and tilted in the coronal plane to advance toward the midline (monkey P, 9°; monkey S, 11°). The same two monkeys were used in previous studies that described different findings on neuronal responses in area V4 (*9, 31*). All experimental procedures were approved by the Institutional Animal Care and Use Committee of the University of Chicago and followed the U.S. National Institutes of Health guidelines.

#### Behavioral task

Monkeys sat in a primate chair facing a calibrated CRT display (1024 × 768 pixels, 100 Hz refresh rate) at 57 cm viewing distance inside a darkened room. Binocular eye position and pupil area were recorded at 500 Hz using an infrared camera (EyeLink 1000, SR Research). Trials started once the animal fixated within 1.5° of a central white spot (0.1° square) presented on a mid-level gray background (**Fig. 1A**). The animal had to maintain fixation until its saccade response at the end of the trial. After a randomly selected fixation period of 400 to 800 ms, two achromatic Gabor sample stimuli appeared for 200 ms, one in each visual hemifield. After a random variable delay of 200 to 300 ms, a Gabor test stimulus appeared for 200 ms at one of the two stimulus locations, selected randomly with equal probability. Shortly after the test stimulus turned off (100 to 150 ms), two saccade targets of different color and shape appeared in opposite directions along an imaginary line orthogonal to the axis of the samples. A go-signal (fixation spot turn off) that occurred 150 to 200 ms after the target onset indicated the animal should make a saccade to the appropriate saccade target according to Gabor orientation change. The locations of the saccade targets were switched randomly across trials. This eliminated motor criterion from the attention-related behavioral *d'* and perceptual decision. Spatially selective attention between the two locations was alternated in blocks of trials (120 to 220). Attention was controlled using different reward size for the two sides, for hits and correct rejections, or the probability of test trials. The location of attention was cued by few instruction trials with a single sample stimulus (10 to 15) before each block started. Five different orientation changes (10° to 90°) were used on the valid side (cued by instruction trials) and one intermediate orientation (28°) was used on the invalid side. Attentional effort (**Fig. 4**) was controlled between low and high values in alternate blocks by switching reward size from small to large, but remained equal between the two locations (*9*). Behavioral task was controlled using custom-written software, LabLib (https://github.com/MaunsellLab/Lablib-Public-05-July-2016.git).

#### Neurophysiological recordings

Extracellular neuronal signals from the multielectrode linear array (16 channel V-probes with 2 optic fiber channels; Plexon Inc.) were amplified, band-pass filtered (250 to 7,500 Hz), and sampled at 30 kHz using a data acquisition system (Cerebus, Blackrock Microsystems) (**Figs. 1** and **2**). We simultaneously recorded from multiple single units and small multiunits over 51 sessions (29 sessions for monkey S; 22 sessions for monkey P). Spikes from each electrode were sorted offline (Offline Sorter, Plexon Inc.) by manually well-defining cluster boundaries using principal components analysis and waveform features. Well-isolated clusters were classified as single units from multiunits based on the isolation quality of unit clusters. The degree to which unit clusters were separated in two-dimensional (2D) spaces of waveform features (first three principal components: peak, valley, and energy) was measured by multivariate analysis of variance (MANOVA) *F* statistic using Plexon Offline Sorter (Plexon Inc.). A unit cluster of MANOVA $P < 0.05$ was considered as a single unit that indicates that the unit cluster has a statistically different location in 2D space and that the cluster is statistically well separated.

<u>Viral vectors</u>

Excitatory opsins, channelrhodopsins (hChR2) were unilaterally expressed in NE-expressing neurons in the LC using a two-virus strategy (*12*). The first vector delivered Cre-recombinase under the control of the dopamine b-hydroxylase (DBH; plasmid pEMS1113, catalog# 29042, Addgene) promoter (AAV9-DBH-Cre-P2A-mCherry-SV40pA, titer of $2.18 \times 10^{13}$ vg/ml; Lot#17-502, Virovek, Inc.). The second virus (AAV5-EF1a-DIO-hChR2(H134R)-EYFP, titer of $2.7 \times 10^{13}$ vg/ml; SKU# VB4652, Vector Biolabs) delivered a Cre-recombinase-dependent hChR2 construct. A similar approach has been successfully used to target midbrain dopaminergic neurons in rhesus monkey (*12*).

<u>Virus injection</u>

Injection sites were identified by electrophysiological recording of LC neurons characterized by waveform shape, sensitivity to arousing brief auditory noise stimuli, and was guided by neuronal response properties of other brain areas along the recording trajectories leading to the LC, which included the superior colliculus, inferior colliculus, and the trochlear decussation in the brainstem (*13, 14*). We injected a volume of ~40 µl and ~50 µl of the two virus combination (1:1; titer $1 \times 10^{12}$ vg/ml) respectively in the monkey S and monkey P in three separate unilateral locations in and around LC using custom-built injection cannulas. The injection cannula incorporated a glass pipette, stainless steel tubing with an insulated tungsten microwire for simultaneous neurophysiological recording and microinjection. We filled the injection cannula with the virus combination (~60 µl) and a calibrated glass tubing provided for visual confirmation of fluid injection. The virus-filled cannula was connected to a pneumatic pump (PV820, World Precision Instruments, customized to incorporate a constant holding pressure) using a patch-clamp microelectrode holder (AM Systems) for the microinjections. We injected at a rate of ~50 nl/min using a constant holding pressure. Behavior and electrophysiology started after 10 weeks after the injection.

<u>Optogenetic stimulation of LC-NE neurons</u>

We used a 120 mW, 457 nm laser (Laserglow Technologies) with power at the end of the optic fiber in the range of ~50 mW. The power and timing were regulated by a Pockels cell (ConOptics) through the same software used for behavioral task (LabLib). Laser light was delivered either as brief trains (e.g., 400 ms train of 10 ms pulses at 20 Hz for behavioral experiments) or 100 ms continuous pulse (for characterizing light-evoked excitability). For optogenetic stimulation sessions (**Figs. 3** and **4**), we used custom made optrodes containing a 100 µm diameter optic fiber (NA, 0.66; Doric Lenses Inc.) attached to a tungsten sharp electrode (FHC Inc.).

<u>Immunostaining</u>

We have performed histological confirmation in one of the experimental monkeys (monkey S) as we are still conducting additional experiments on monkey P. First, the animal was deeply anesthetized and then euthanized, and then perfused with phosphate buffer solution (PBS) followed by 4% paraformaldehyde in PBS. The brain was removed, cryoprotected (in graded sucrose solutions, 10, 20 and 30%) and blocked. Sections of 50 μm were cut using cryostat and stored in PBS. Then the sections were washed in PBS containing 1% triton (1% PBS-T) for 30 min followed by washing in PBS containing 0.3% triton (0.3% PBS-T) for 5 min. Then sections were placed sequentially in blocking buffer (5% goat serum in 0.3% PBS-T, Cat# 01-6201, ThermoFisher) for 3 h and in primary antibody solution (DBH antibody, 1:500, Cat# 22806, Immunostar; mCherry Monoclonal Antibody, 1:2000, Cat# 16D7, ThermoFisher; GFP Monoclonal Antibody, 1:1000, Cat# GF28R, ThermoFisher; in blocking buffer) for 48 h in 4ºC. Then sections were washed in 0.3% PBS-T and incubated in secondary antibody solution (Goat anti-Rabbit IgG (H+L) Highly Cross-Adsorbed Secondary Antibody, Alexa Fluor 647, 1:500; Goat anti-Rat IgG (H+L) Cross-Adsorbed Secondary Antibody, Alexa Fluor™ 594, 1:500; Goat anti-Mouse IgG (H+L) Cross-Adsorbed Secondary Antibody, Alexa Fluor™ 514, 1:500; ThermoFisher, in 0.3% PBS-T) for 4 h in room temperature. Following sequential washes in 0.3% PBS-T and PBS, sections were mounted with DAPI mounting media (Cat# P36981, ThermoFisher). Two days later sections were imaged with confocal microscope.

### Data analysis

#### *Behavioral analysis*

All completed trials were included in our analysis. Behavioral sensitivity ($d'$) and criterion ($c$) at a spatial location were measured from hit rates within nonmatch trials and false alarm (FA) rates within match trials using 1-dimensional signal detection theory (*31, 32*) as: $d' = \Phi^{-1}(H) - \Phi^{-1}(F)$ and $c = -\frac{1}{2}[\Phi^{-1}(H) + \Phi^{-1}(F)]$; where $\Phi^{-1}$ is inverse normal cumulative distribution function; H and F are respectively the rates of hits and false alarms.

#### *Pupil area*

All pupil area measurements were measured binocularly at 500 Hz while monkeys maintained fixation in the absence of a luminosity change using infrared camera (EyeLink 1000, SR Research). Raw pupil areas were *z* scored for each session and each eye separately. Mean pupil area was measured by averaging the *z*-scored pupil area during the 400 ms after sample appearance.

#### *Neuronal response*

Spike counts in 2 ms bins were smoothed using half-Gaussian kernel (standard deviation of 15 ms, rightward tail) for PSTHs. Neuronal responses were classified according to their average responses (greater than 250 ms pre-event period values at least over consecutive 20 bins; $p < 0.05$) to sample stimulus (over 50 to 300 ms from sample onset; phasic), saccade (over –50 to 200 ms from saccade onset; saccade responsive) and during fixation period (over 0 to 400 ms from fixation; non-phasic). Spike trains were converted into z-scores (normalized with respect to mean and standard deviation of spike rates over 250 ms pre-event duration) to construct population PSTHs. Neuronal modulation with spatial attention was measures by neuronal sensitivity as: $d'_{neuron} = (\mu_{contra} - \mu_{ipsi}) / \sqrt{\frac{1}{2}(\sigma_{contra}^2 + \sigma_{ipsi}^2)}$; where $\mu_i$ and $\sigma_i$ are the average and standard deviation of spike counts within 50 to 300 ms and –400 to 0 ms from sample stimuli onset respectively for phasic and non-phasic neurons (*i*, attended location, contra- or ipsi- lateral to the recorded LC hemisphere).

#### *Demixed principal component analysis*

We examined how well LC neuronal activity during the presentation of sample stimuli could decode the focus of selectivite attention, and whether the response on a trial would be correct.

These parameters were isolated from other stimuli and task variables using demixed principal component decompositions (*11*) (**figs. S5, S6; Fig. 2**). All responsive LC units from both monkeys were separated into phasic and non-phasic neurons. Details of the demixed principal component decompositions have been described by Kobak *et al.* (*11*). Briefly, neuronal population activity patterns were decomposed into a linear combination of specific components, each of which carries information of a single task variable. Mean-subtracted and trial-averaged spike trains of each neuron were decomposed into sum of marginalized averages, each corresponding to a task variable and a noise term. This marginalization process ensures that the individual components are uncorrelated. A loss function that penalizes the difference between the marginalized data and the reconstructed full data is minimized using the least-square method. The reconstructed data are the full data projected with the decoders onto a low-dimensional latent space and then reconstructed with the encoders. Such decomposition method offers reliable decoding accuracy measures even for relatively small explained variances by indivual variables (**figs. S5, S6**) (*11*). Trials were classified into two attention conditions (attend contra versus ipsi) and two stimulus detection conditions (correct (hit, CR) versus error (miss, FA)), yielding four different trial configurations (**Fig. 2A, 2C**). Single-trial spike rates were filtered with a half Gaussian kernel ($\sigma$ = 30 ms) and subsampled at 100 Hz. We analyzed spike rates over 500 ms (50 time points, starting 100 ms before sample onset) for phasic responsive units and over 1000 ms (100 time points, starting 800 ms before sample onset) for non-phasic responsive units. Decomposition into demixed components was performed on training datasets (leave-one-out, 1000 repetitions). Attended location and detection were then decoded on the remaining cross-validated test trials using the top three components to estimate the decoding accuracy (**Fig. 2B, 2D**).

*Statistical analysis*

Unless otherwise specified, we used paired *t* test and multifactor ANOVA for comparing normally distributed datasets. Normality was checked using a Kruskal-Wallis test.

**Supplementary figures**

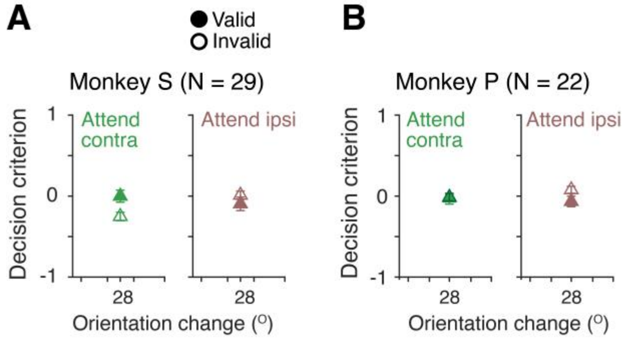


**Fig. S1. Session averaged behavioral decision criteria associated with sptatially selective attention.** (**A-B**) Average behavioral criteria for valid (filled triangles) and invalid (open triangles) trials when monkeys' attention was directed to the location either contra- or ipsi- lateral to the recorded LC. Error bars, 95% confidence intervals.

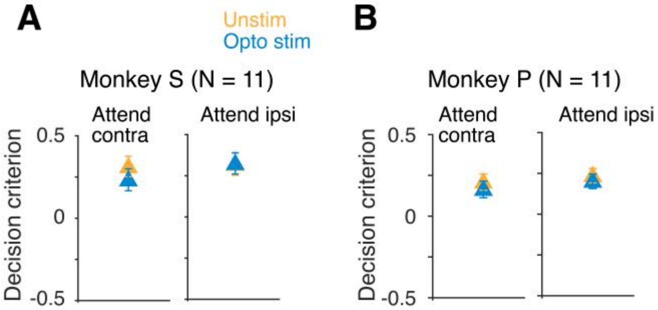


**Fig. S2. Effects of optogenetic stimulation on behavioral decision criteria associated with spatially selective attention.** (**A-B**) Average behavioral criteria for all valid trials separated according to optogenetically stimulated and unstimulated trials for the same dataset shown in **Figs. 3E-F**. Trials are also grouped according to whether monkeys' spatially selective attention was directed at the location either contra- or ipsi- lateral to the recorded LC. Error bars, 95% confidence intervals.

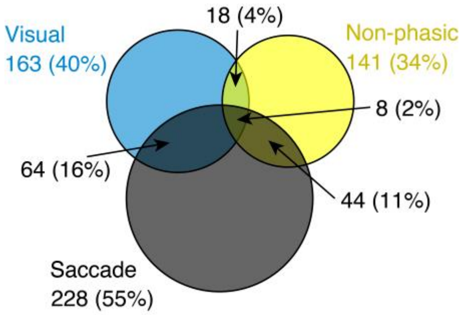


**Fig. S3. Proportions of different neuron types in the LC according to their task-specific responses.** Venn diagram of phasic (visual), non-phasic and saccade responsive units in the LC from both experimental monkeys. Total, 414 units (monkey S, 229; monkey P, 185); visual, 163 (monkey S, 89/229; monkey P, 74/185); non-phasic, 141 (monkey S, 88/229; monkey P, 53/185) and saccade responsive, 228 (monkey S, 104/229; monkey P, 124/185).

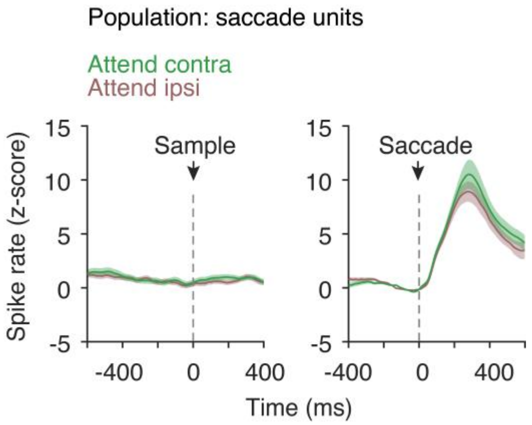


**Fig. S4. PSTHs of saccade responsive units in the LC.** Population averaged spike rates of all saccade responsive LC neurons aligned with respect to sample stimuli onset (*left*) and saccade onset (*right*). n = 228 (monkey S, 104; monkey P, 124). Error bars, ± 1 SEM.

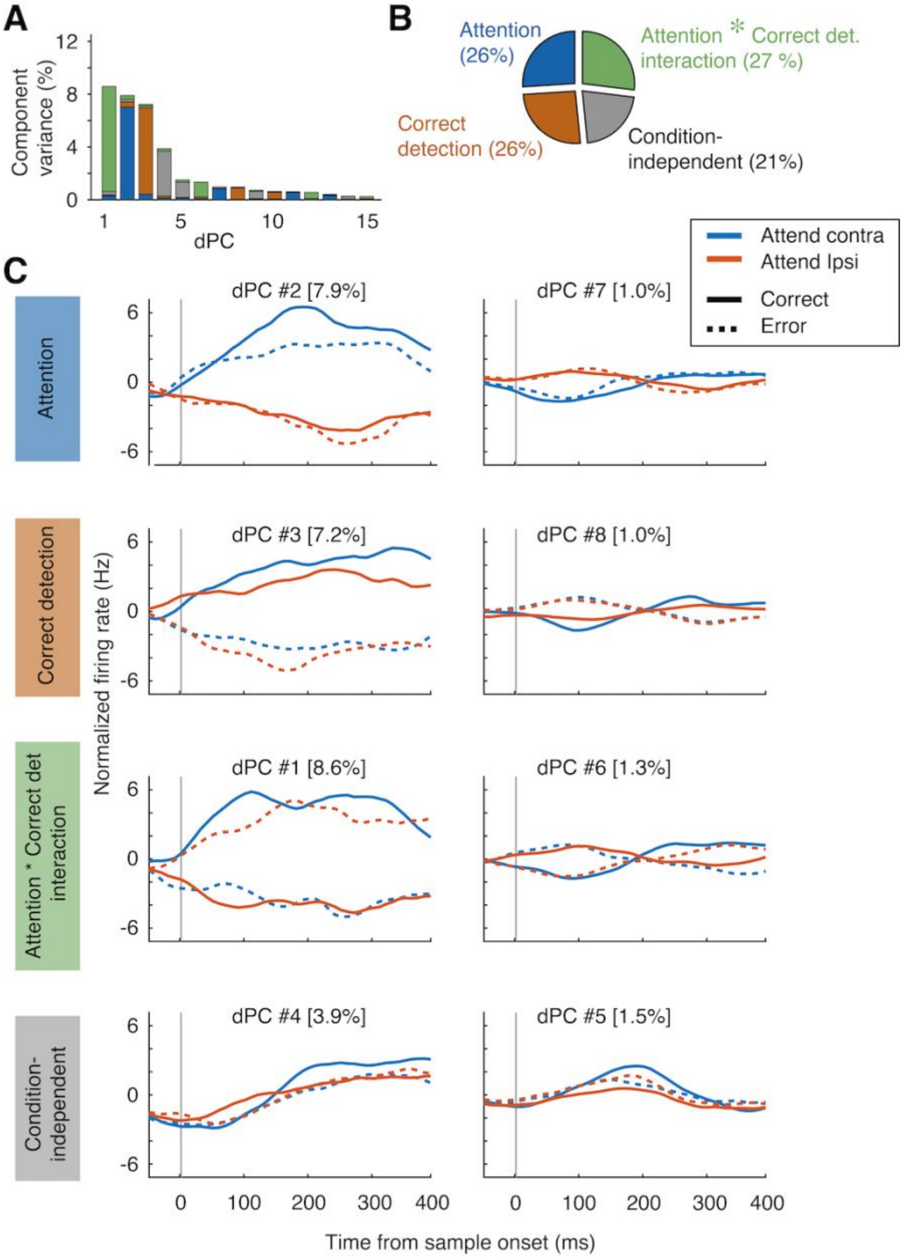


**Fig. S5. Demixed principal component decomposition of population spike rates of phasic responsive LC neurons.** Population averaged spike rates for all phasic responsive LC neurons were decomposed into components by systematically capturing most of the variance of the data considering two experimental conditions: the location of attention (contra versus ipsi to the recorded LC) and behavioral detection of orientation changes (correct versus error trials). Information about these variables has been isolated from other task variables. Single-trial spike rates were filtered with a half Gaussian kernel (σ = 30 ms, right tail) and subsamples at 100 Hz. Spike rates over 500 ms (50 time points) starting from 100 ms before the sample stimulus onset were used for the analysis. (**A-B**) Total explained variance (percent) for individual demixed principal components corresponding to different task variables (A) and their collective proportions ((B), first 20 components). (**C**) Demixed principal components arranged according to the task conditions. Explained variance of each component is shown. Traces in each subplot are the projections of spike rate PSTH data onto the respective demixed principal component decoder axis and correspond to 4 task conditions. *Top row*, first two components for selective attention. *Second row*, first two components for perceptual detection. *Third row,* first wo components for the interaction between attention and detection. *Forth row*, attention and detection independent components.

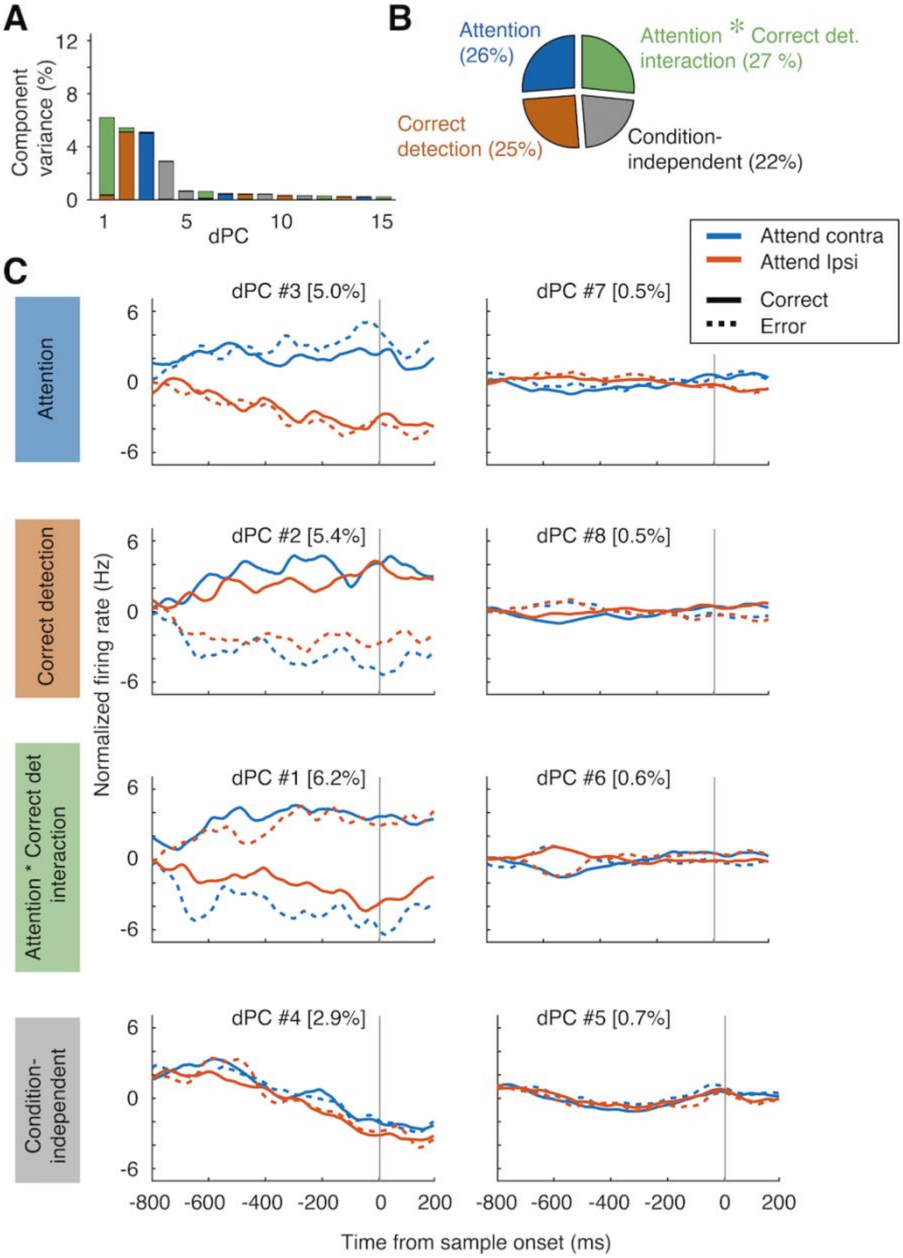


**Fig. S6. Demixed principal component decomposition of population spike rates of non-phasic responsive LC neurons.** Demixed principal components from population of non-phasic responsive LC neurons same as in **Fig. S5.** Single-trial spike rates were filtered with a half Gaussian kernel (σ = 30 ms, right tail) and subsamples at 100 Hz. Spike rates over 1000 ms (100 time points) starting from 800 ms before the sample stimulus onset were used for the analysis. (**A-B**) Total explained variance for individual demixed principal components corresponding to different task variables (A) and their collective proportions ((B), first 20 components). (**C**) Demixed principal components arranged according to the task conditions. Explained variance of each component is shown. Traces in each subplot are the projections of spike rate PSTH data onto the respective demixed principal component decoder axis and correspond to 4 task conditions. *Top row*, first two components for selective attention. *Second row*, first two components for perceptual detection. *Third row*, first two components for the interaction between attention and detection. *Forth row*, attention and detection independent components.

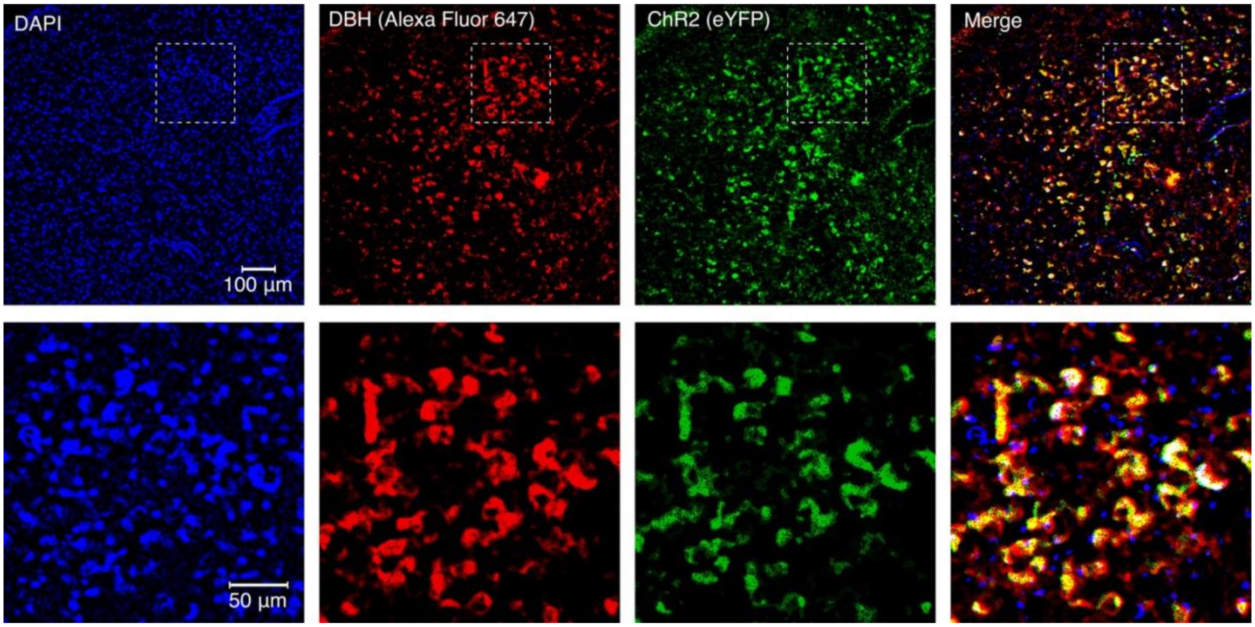


**Fig. S7. Immuno-histological confirmation of ChR2 expression in LC-NE neurons in monkey**. A representative immuno-stained coronal brain section of monkey S showing colocalization of NE neurons and ChR2 in the LC. *Panels from left to right,* DAPI nuclear staining, Dopamine-beta-Hydroxylase (DBH) immune-staining of LC-NE neurons, Enhanced yellow fluorescent protein (eYFP) in DBH:Cre:hChR2 (AAV5-EF1a-DIO-hChRh2(H134R)-EYFP) neurons in the LC, and merged image of DAPI, DBH and eYFP. Yellow color cells represent colocalization of eYFP and DBH. Bottom panels represent respective high magnification images of the box area in the top panel.
